# TransporterPAL: An integrative database Transporter Prediction ALgorithm

**DOI:** 10.1101/2022.09.05.506577

**Authors:** Jane Dannow Dyekjær, Alexander Kruhøffer Bloch Andersen, Joel August Vest Madsen, Jens Preben Morth, Lars Juhl Jensen, Irina Borodina

## Abstract

**Motivation:** Natural products are used as drugs, cosmetic ingredients, pigments, flavors, and agricultural products. The compounds are retrievable as extracts from natural sources, but the yields are often low, and the final product may contain various impurities. These challenges can be solved by expressing the biosynthetic pathway in microbial cell factories and ensuring product secretion from the cell by using an appropriate transporter. However, insufficient knowledge of transporters for specific compounds often obstructs efficient secretion of the natural product. Therefore, our goal was to develop an algorithm that predicts transporters for a given compound using available public data.

**Results:** The web application TransporterPAL predicts suitable transporters for compounds by interconnecting data for biosynthetic genes and their interactions with transporters. The web application queries the STITCH, STRING, and UniProtKB databases via their respective APIs and returns a set of potential transporters based on a compound and, optionally, the organism as input. For a test set of 61 transporter systems, each containing one or more transporters, a total of 90 unique transporters with a known substrate, we could retrieve 45% of the transporters. To our knowledge, this is the first bioinformatics tool for predicting transporter candidates for a given molecule.

**Availability:** https://transporterpal.com

**Supplementary information:** Supplementary data are available at Bioinformatics online.

## 1 Introduction

The demand for natural products is increasing in the pharmaceutical and biotechnological industries. Due to the complexity of these molecules, the chemical synthesis of natural products is usually challenging, if not impossible. Alternatively, natural products can be produced cheaper and more sustainably by incorporating heterologous pathway genes into microbial cell factories. Secretion of the synthesized molecules from microbial cells is vital to alleviate cellular toxicity, prevent product degradation, and allow for high titer and purity (van der Hoek and Borodina, 2020). Often, a specific transmembrane transporter protein is required to engineer the secretion of small molecules. However, transmembrane proteins are challenging to study, and many transporters are thus not well characterized. As a result, the knowledge of appropriate transporters is scarce, but the exploitation of data from several independent databases can provide clues about potential transporters.

We hypothesized that if we can extract pathway genes for biosynthesis of a compound and other proteins interacting with the compound, we can identify a suitable transporter among their interaction partners in a functional association network. Furthermore, assuming the transporter is present within this expanded set of proteins, we can use text mining to differentiate transporters from other pathway proteins based on their functional description. Here, we present a web resource that interacts with public databases to predict potential transporters automatically for a given compound produced in nature, irrespective of protein annotation.

## 2 Implementation

STITCH (Szklarczyk *et al*., 2016) is a database containing information about chemical–protein interactions from many sources, including biological pathway databases, automatic text mining of biomedical literature, and experimental data repositories. The STRING database (Szklarczyk *et al*., 2021) similarly provides physical and functional protein-protein interactions integrated from various sources. Finally, UniProtKB (Bateman *et al*., 2021) provides functional annotations of proteins, including Gene Ontology (GO) annotations related to transport activity.

Our workflow consists of the following steps, starting from a chemical and, optionally, an organism of interest: (i) use chemical–protein interactions from STITCH to obtain an initial set of proteins (Szklarczyk *et al*., 2016), (ii) expand this set of proteins with their interaction partners using functional associations from STRING (Szklarczyk *et al*., 2021), (iii) select the transport-related proteins from this set based on UniProtKB GO annotations (Figure 1). To combine the information from all these databases, we access them via their respective REST APIs. Specifically, we use the *interactors* API method of STITCH to obtain interacting proteins with the input chemical, then the *interaction_partners* API method of STRING to expand the set of proteins. We convert the STRING identifiers to UniProt IDs using the UniProt Retrieve/ID mapping service and subsequently retrieve the UniProtKB record for each ID via the UniProt API. Each record contains information about the protein, and as we search specifically for transporters, we filter the records based on keywords related to transport activity. We have used UniProt GO annotations for cellular components, molecular functions, and biological processes to generate a list of transporter-related keywords. After removing protein entries unrelated to transport, the UniProtKB accession number, protein name, and organism name are retrieved from UniProtKB. The interaction between the compound and each putative transporter is scored using the combined score from the network STITCH API, as described in (Szklarczyk *et al*., 2016).

**Figure 1.**
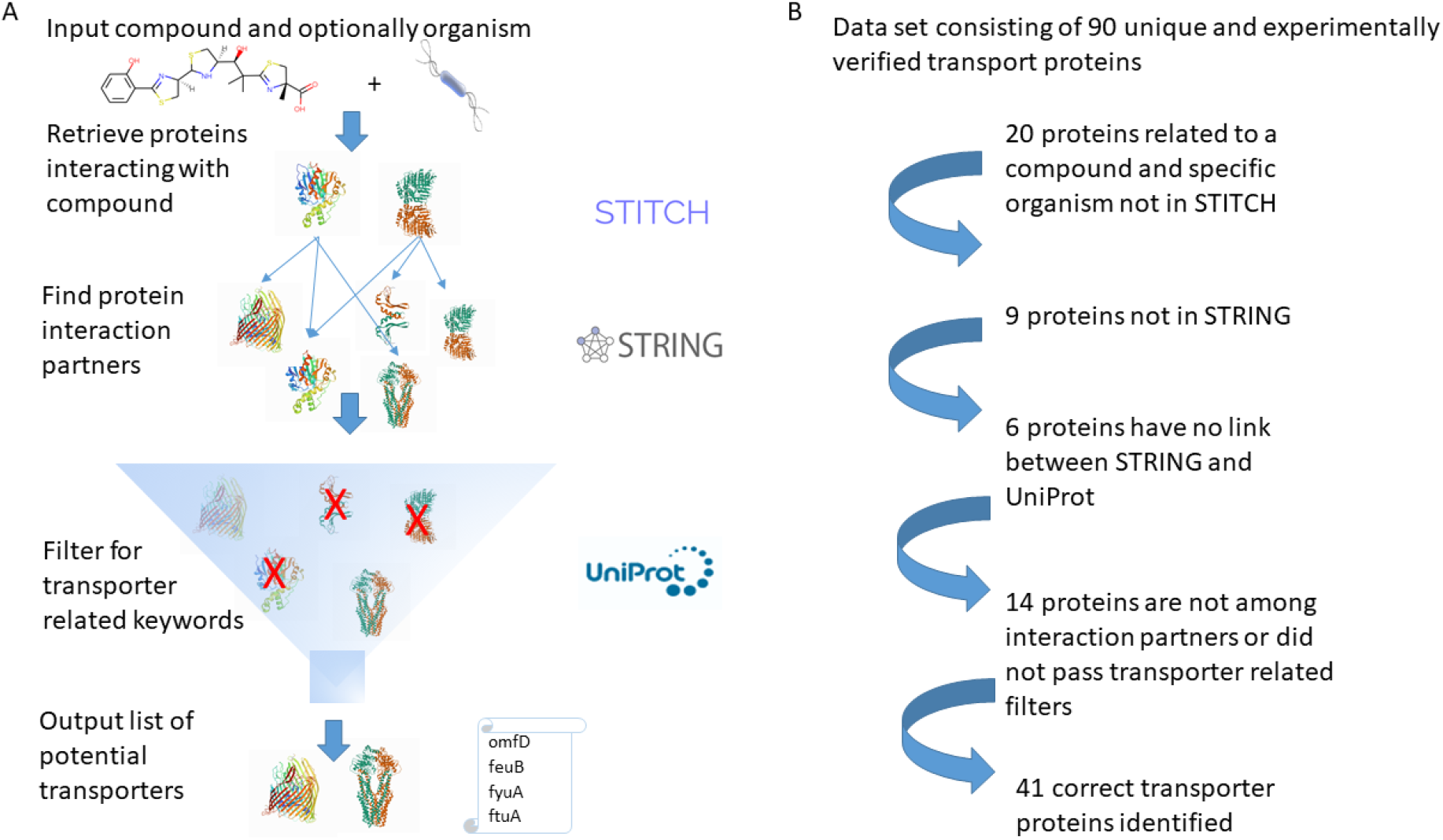
TransporterPAL workflow for transporter identification. Panel A. We use the STITCH database to extract pathway genes and proteins interacting with the compound for an organism. This set of proteins is expanded by their interaction partners using STRING. The list contains non-transporter-related proteins, and we exclude these by filtering for transporter-related GO annotations using UniProt to obtain a final list of potential transporters for the compound. Panel B. 90 unique, experimentally verified transporter proteins were used in the test set. Twenty proteins related to a compound and specific organism were unavailable in STITCH. Nine proteins were not present in the STRING, and six transporters did not have a link between the STRING and UniProt databases. In the remaining data set of 55 transporters, 14 were not found, resulting in 41 correctly identified transporter proteins.

The web application is based on Python3 and uses STITCH, STRING, and UniProt APIs. We use the python packages requests, csv, json and sys. Furthermore, asyncio and aiohttp are used to make asynchronous API calls. The web application has been developed using an Express NodeJS web application framework in combination with the basic web application building blocks, HTML, CSS, and Javascript.

## 3 Availability

On the online server https://transporterpal.com, the user enters a natural compound and, optionally, an organism. A link to available organisms is provided on the website. When the web application has finished, it sends an email containing a CSV file and a FASTA file containing the amino acid sequences of the identified transporters.

## 4 Current benchmarking

Although the algorithm works with any organism, we tested the algorithm on a data set of bacterial transporters extracted from the Transporter Classification Database (Saier *et al*., 2021), a database containing experimentally verified transporters, to quantify the number of transporters we can retrieve by using the algorithm. We chose transporters of degradation pathways and secondary metabolite biosynthesis produced by specific bacteria and corroborated their pathway in MetaCyc (Caspi *et al*., 2020). We selected transporters from 61 transporter classification IDs, each containing one or more transporters, in total 95 transporters (Supplementary data). A few of the transporters mediate transport of more than one compound, which makes 90 of the transporters unique. Of the unique transporters, 35 were not present in the databases; For 20 compounds, transporters related to a specific organism were unavailable in STITCH, primarily due to unrepresented species. Additional nine transporters were not present in the STRING database. Furthermore, six transporters did not have a link between the STRING and UniProtKB databases. From the remaining data set of 55 proteins, 14 transporters were not found either because they were not among the interaction partner proteins or because they did not contain the transporter-related GO-annotation keywords. Ultimately, 41 proteins were correctly retrieved, corresponding to 46% of the 90 unique transporters. The predictive power of TransporterPAL depends on available data, highlighting the need for more experimental research on transporter function characterization.

## 5 Conclusion

We have developed a web application to predict potential transporters based on the pathway genes and interacting proteins of a given compound and organism. With more research on transporters and the addition of knowledge into the STITCH, STRING, and UniProtKB databases, we expect to increase the success rate of retrieving suitable proteins with transport activity for a specific compound. While we focus on transporters, this method is easily extendable to study other pathway proteins by changing the filters in the GO annotation to keywords suitable for a particular protein family class. To our knowledge, this is the first tool able to link potential transporters to a given natural product.

